# Reactive oxygen species signalling is involved in alkamide-induced alterations in root development

**DOI:** 10.1101/2021.12.23.474045

**Authors:** Tonatiu Campos-García, Jorge Molina-Torres, Kirk Overmyer

**Author notes:** Corresponding Author: Kirk Overmyer; University of Helsinki, P.O Box 65 (Viikinkaari 1), FI-00014 Helsinki, Finland.

## Abstract

Alkamides are alpha unsaturated *N-*acylamides structurally related to *N*-acyl ethanolamides (NAEs) and *N*-acyl-L-homoserine lactones (AHLs). Studies have shown that alkamides induce prominent changes in root architecture, a significant metabolic readjustment, and transcriptional reprogramming. Some alkamide responses have been associated with redox signalling; however, this involvement and ROS sources have not been fully described. We utilized a genetic approach to address ROS signalling in alkamide-induced processes and found that in *Arabidopsis*, treatment with the alkamide affinin (50µM) increased the *in-situ* accumulation of H_2_O_2_ in lateral root emergence sites and reduced H_2_O_2_ accumulation in primary root meristems implying that altered root growth was dependent on endogenous H_2_O_2_. Results show that ROS sourced from PRX34, RBOHC and RBOHD were involved in promotion of lateral root emergence by alkamides. RBOHC was required for affinin-induced enhanced root hair expansion. Furthermore, affinin-induced changes in lateral root emergence, but not root hair length, were dependent on a change in extracellular pH. Finally, reverse genetic experiments suggest heterotrimeric G-proteins were involved in plant response to alkamides; nevertheless, further studies with additional higher order G-protein mutants will be required to resolve this question. These results support that alkamides recruit specific ROS signaling programs to mediate alterations in root architecture.

**Highlight:** Reactive oxygen species (ROS) are involved in alkamide-induced altered root development. Heterotrimeric G-protein complex, extracellular acidification, and ROS sourced from peroxidases and NADPH-oxidases are involved in these processes.

## Introduction

In plants and animals some metabolites interact with messengers or receptors to modulate diverse metabolic pathways and signal transduction cascades. Plant chemical effectors from different sources can trigger physiological and morphological responses. Alkamides are low molecular weight α-unsaturated acyl-amide plant metabolites distributed in some plant species. These metabolites are known to have a wide range of biological activities in bacteria, fungi, plants, and mammals (Molina-Torres *et* al. 1996; Gertsch, 2008; Prachayasittukal *et al*. 2013). These include antimicrobial and other activities that suggest these compounds may function in defence against competing plants, microbes, and herbivorous pests (Molina-Torres *et al*. 2004) Affinin (*N*-isobutil-*2E,6Z,8E*-decatrienamide) is the most abundant alkamide found in the ethanolic root extracts from chilcuague [*Heliopsis longipes* (A. Gray) S. F. Blake], an endemic species from “Sierra Gorda”, México. This molecule has been shown to have bacteriostatic and fungistatic effects (Molina-Torres *et al*. 1996) and to alter plant root growth and development (Ramírez-Chávez *et al*. 2004). More detailed plant developmental studies have found that alkamides also alter signalling in plants, modulating both developmental and stress response pathways, functioning as biochemical elicitors (López-Bucio *et al*. 2007; Méndez-Bravo *et al*. 2011). Most organisms contain amide lipids composed of one or two amines linked to a fatty acid through an amide bond in their inner and outer membranes. *N*-acylethanolamides (NAEs), which include some endocannabinoids, are an example of amine lipids structurally related to alkamides that have important biological signalling functions in plants (Blancaflor *et al*. 2003; Coulon *et al*. 2012; Blancaflor *et al*. 2014) and in mammals (Kunos *et al*. 2000; Wilson and Nicoll, 2002). Faure *et al*. (2014) found that alkamides could alter the NAE metabolic pathway by modulating fatty acid amide hydrolase (FAAH) activity. However, they also concluded that metabolic pathways other than FAAH are also involved in the metabolism of alkamides in plants, indicating the need for further studies in this area. Alkamides also exhibit structural similarities to *N*-acyl-homoserine lactones (AHLs), another amino lipid compound used by many bacteria as quorum-sensing signals to coordinate their collective behaviour. Accumulating evidence indicates that AHL are also perceived by plants, which respond by altering cell immune responses, host defence, stress responses, energy and metabolic activities, transcriptional regulation, protein processing, cytoskeletal activities, root development and plant hormone responses (Mathesius et al. 2003; Kravchenko et al. 2006).

Affinin isolated from *Heliopsis longipes* roots displays a dramatic effect on *Arabidopsis thaliana* (*Arabidopsis*) root system architecture by altering primary root growth, lateral root emergence, and increasing root hair elongation in a dose dependent manner (Ramírez-Chávez *et al*. 2004). Méndez-Bravo *et al*. (2010) found that *N*-isobutyl decanamide and the interacting signals JA and nitric oxide (NO) act downstream independently of auxin-responsive gene expression to promote lateral root formation and emergence, providing compelling evidence that NO is an intermediate in alkamide signalling mediating the root system architecture alterations observed in *Arabidopsis*. Transcriptomic profiling of *Arabidopsis* has shown that exogenous application of *N*-isobutyl decanamide triggers profound physiological changes with activation of developmental, defence, and stress related genes (Mendez-Bravo *et al*. 2011). Transcripts of several JA-related genes such as *PDFs, VSP2, JAZ10* and *JAZ8* accumulated to higher levels. Transcripts of at least 70 genes belonging to the functional group ‘‘oxygen and radical detoxification’’, also exhibited enhanced accumulation by *N-*isobutyl decanamide treatment. They found that alkamides could modulate some - defence responses associated with necrotrophic pathogens through JA-dependent and MPK6-regulated signalling pathways. Additionally, decanamide induces ROS accumulation in leaf tissue (Mendez-Bravo *et al*. 2011). These results suggest that general defence-associated responses elicited by *N*-isobutyl decanamide and affinin appear to be related to both hormone and ROS signalling pathways.

Recent studies revealed that ROS act as essential signalling molecules in plants and are required for several basic biological processes including cellular proliferation and differentiation, immunity, and stress responses (Mittler, 2017). ROS signalling specificity is dependent on the species produced and their subcellular location. It is well known that ROS, such as superoxide anion (O_2_^•-^), hydrogen peroxide (H_2_O_2_), hydroxyl radical (^•^OH), and singlet oxygen (^1^O_2_), play important signalling roles in plants as key regulators of growth, development, response to biotic and environmental stimuli, plant metabolism, and programmed cell death (Jaspers and Kangasjärvi, 2010; Suzuki *et al*., 2011; del Río, 2015; Overmyer et al., 2003). In plants, H_2_O_2_ is the product of catalytic reactions from different enzymes such as the peroxisomal flavin-containing enzymes glycolate oxidase and the acyl-CoA oxidase, which are involved in the photorespiratory and fatty acid β-oxidation pathways, respectively. H_2_O_2_ is produced in the chloroplast under normal and stress conditions, as well as from the activities of peroxidases, and in some species, oxalate oxidase (Torres, 2010; Baxter et al., 2014). It is now well established that a major source of the O_2_^•-^ is the plasma membrane-localized NADPH-oxidases (del Río, 2015). O_2_^•^ is a charged molecule under most physiological conditions and can not passively transfer across a membrane. However, O_2_^•-^ can dismutate into H_2_O_2,_ either enzymatically by superoxide dismutase (SOD), peroxidases or spontaneously, especially at low pH. H_2_O_2_ can readily cross membranes passively or via aquaporins. Additionally, O_2_^•-^ can mediate the formation of membrane soluble lipid peroxides (Miller *et al*., 2010; Mittler *et al*., 2011). In many biological systems (as animals and plants) ROS signalling is mediated by a highly regulated process of ROS accumulation in specific cellular compartments. The NADPH-oxidases, termed *NOX* in animals and RESPIRATORY BURST OXIDASE HOMOLOGs (RBOHs) in plants, are plasma membrane localized enzymes that produce ROS into the apoplast. This family of enzymes are highly regulated via calcium and various phosphorylation/dephosphorylation events (Ogasawara *et al*. 2008; Mittler, 2017). In *Arabidopsis* the ROS produced by the NADPH-oxidases RBOHC, RBOHD, and RBOHF, as well as the class III peroxidases, have been shown to be involved in the lateral root emergence and root hair growth (Ogsawara *et al*. 2008; Li *et al*. 2015; Orman-Ligeza *et al*. 2016; Manzano *et al*. 2014, Fernández-Marcos *et al*. 2017). Nevertheless, the source of alkamide-induced ROS production and the signalling pathways recruited downstream that lead to modifications in root growth, development, and plant metabolism remain unknown. Here we utilize a genetic approach to address the role of ROS signalling in alkamide-induced processes.

## Materials and Methods

### Plant material and growth conditions

*Arabidopsis thaliana* (*Arabidopsis*) accession Columbia-0 (Col-0) was used for all experiments unless otherwise indicated. For reverse genetic experiments, the following mutants were used; the NADPH-oxidases, *respiratory burst oxidase homologC* (*rbohC), rbohD*, and *rbohF*; class III peroxidase, *peroxidase34* (*prx34*); heterotrimeric G-protein subunits, g*-protein alpha subunit1-2* (*gpa1*-2*), arabidopsis gtp-binding protein beta1*-2 (*agb1*-2*), arabidopsis g-protein gamma subunit2*-1 *(agg2*-1), and the *regulator of g-protein signaling1*-2 (*rgs1*-2). Mutant seeds were obtained from NASC (www.arabidopsis.info). Seeds were surface sterilized with 70% (v/v) ethanol plus 2% triton-X 100 for 5 min, then rinsed in absolute ethanol for 1 min, and dried in a laminar hood over sterile filter paper sheets. Seeds were germinated and grown on agar plates (9 g L^-1^) containing 0.3x MS medium and sucrose (11 g L^-1^). Seeds were stratified 72 hrs in darkness at 4°C and then placed in controlled environment growth chambers (Weiss FITOTRON SCG120; www.fitotron.co.uk) vertically to allow root growth along the agar surface and unimpeded hypocotyl growth. Plants were grown under a long-day photoperiod (16 h light, 8 h darkness), 25°C/18°C day/night temperature and a light intensity of 60-100 µM m^-2^ s^-1^. After germination for four days, seedlings were transferred to plates containing solvent control or affinin treatments (10 or 50 µM). For the pharmacological assay, one-week old seedlings were transferred to a solid medium supplemented individually or in various combinations of affinin (7 or 50 µM), diphenyleneiododium (DPI; 0.3μM), and solvent control for seven days. In MES buffer experiments, four-day-old seedlings were transferred to medium supplemented with MES buffer (0.5g L^-1^); after the addition of all the reagents the pH was adjusted to 5.7 with NaOH and then autoclaved. After autoclaving, treatments where prepared adding the affinin (10 or 50 µM) to the medium. After seven days the primary root length, number of emerged lateral roots and root hair length were evaluated.

### Affinin isolation and purification

Affinin extraction was performed as previously reported by Ramírez-Chavez *et al*. (2004). After extraction, the concentrated oil was weighted, 2 g were purified by column chromatography as previously reported by Ramírez-Chávez *et al*. (2004); subsequently, the purified fraction was analysed by GC/EIMS to confirm its purity and concentration.

### ROS staining

H_2_O_2_ accumulation was monitored *in situ* with 3,3′-diaminobenzidine tetrachloride (DAB; Sigma-Aldrich; www.sigmaaldrich.com) based on the method of (Thordal-Christensen *et al*., 1997). Six-day-old seedlings were immersed in DAB (1 mg ml^-1^) in PBS buffer (10 mM; pH 7.0) plus 0.05% (v/v) tween-20 and placed overnight at room temperature in the dark. O_2_^•-^ was detected based on nitroblue tetrazolium (NBT)-reducing activity. Seedlings were covered in an NBT solution (0.5mg ml^-1^) in 0.1M PBS buffer (pH 7.4) plus 0.05% triton-X 100 and incubated in the light for 10 minutes. DAB and NBT reactions were stopped by replacing staining solution with fixative / clearing solution of ethanol / acetic acid / glycerol (3:1:1) for 24 hours, mounted on glass slides with a coverslip and visualized with a Leica 2500 microscope (www.leica-microsystems.com).

### Root growth analysis and microscopy

Root growth measurements were performed on images taken from the plates using the SmartRoot plugin on ImageJ (www.imageJ.net; https://smartroot.github.io/). For the analysis of emerged lateral roots, root samples were visualized with a Leica MZ10F stereo microscope equipped with a model DFC490 digital camera attachment (Leica; www.leica-microsystems.com) and lateral roots protruding beyond the epidermal tissue were scored as emerged. Primary root zone organisation of seedlings with or without 50µM affinin treatment were visualised with 5 mg/ml propidium iodide stain and observed in a Carl Zeiss LSM800 confocal laser microscope (https://www.zeiss.com/). Measurements of distance from primary root tip to first root hair bulge, cell length from the elongation/differentiation zone (EZ/DZ) and apical meristem length (AM) were made with ImageJ software (www.imageJ.net). DAB staining images were taken with a Leica 2500 microscope with differential interference contrast (DIC) optics (https://www.leica-microsystems.com/). For each treatment, at least ten plants were analysed. Representative plants for each treatment were photographed using the Nomarski optics on a Leica 2500 microscope. Quantification of DAB staining intensity was done by ImageJ-based quantification described by Béziat (2017). Prior to quantification, the colour mode of light micrographs was converted from RGB to HSB by using ImageJ software and the saturation channel in the HSB stack was selected. Then the intensity of DAB staining was measured in the saturation channel from a same size area in all pictures. An increase of saturation depicts more “pure” colour, while a decrease denotes a more “washed-out” signal (Béziat *et al*., 2017).

### Bromocresol purple pH assay

To test pH changes in the roots, seven-day-old *Arabidopsis* seedlings Col-0 were transferred to a solid medium (as described before) with different affinin treatments. At 24 hours, seedlings were transferred from treatments to a glass slides covered with 1ml of the pH indicator bromocresol purple (www.sigmaaldrich.com; 0.04 g L^-1^) in agarose (0.7%) prepared with distilled water plus CaSO_4_ (0.2mM) (Zandonadi *et al*., 2010). The pH was adjusted to 7.5 with NaOH. After 30 min, glass slides with the seedlings were photographed with a CAMAG TLC Visualizer (www.camag.com). Photographs were analysed with ImageJ software as follows. First, images were converted to an 8-bit format and calibrated with a set of density standards (pH scale bar), a region of interest (ROI) was selected from each pH scale point and the mean grey value of each of those points was recorded and then fitted to a “curve fitting method (linear function)” from the popup menu as described in the ImageJ user manual (https://imagej.nih.gov).

### Statistical analysis

Data from mutant sets were analysed with two-way ANOVA (*n* = 10) and a Tukey’s posthoc test using the Minitab 14 software (www.minitab.com). Different letters are used to indicate significance groups (means that differed significantly *P* ≤ 0.05). For single genotype experiments, data were analysed with one-way ANOVA (*n* = 10) and Tukey’s posthoc test (*P* ≤ 0.05) using InfoStat software (www.infostat.com.ar).

## Results

### Root phenotype of *Arabidopsis Col-0* in response to affinin

To establish the principal affinin-induced phenotypes in seedlings of WT Col-0 *Arabidopsis* under our conditions, root architecture was monitored in response to two affinin concentrations (10 µM and 50 µM), in aseptic cultures on 0.3x MS solid medium. After seven days of growth, the 10 µM treatment resulted in an increased number of emerged lateral roots without a significant decrease on the primary root length (Fig. 1A-C). The 50 µM treatment reduced growth monitored as primary root length and resulted in a significantly increased number of emerged lateral roots and root hair length (Fig. 1A-D). These results show that affinin treatments influenced *Arabidopsis* root system architecture, consistent with the previously reported phenotypes induced by affinin and decanamide (Ramírez-Chávez *et al*., 2004; López-Bucio *et al*., 2007; Méndez-Bravo *et al*., 2010; Méndez-Bravo *et al*., 2011).

**Figure. 1.**
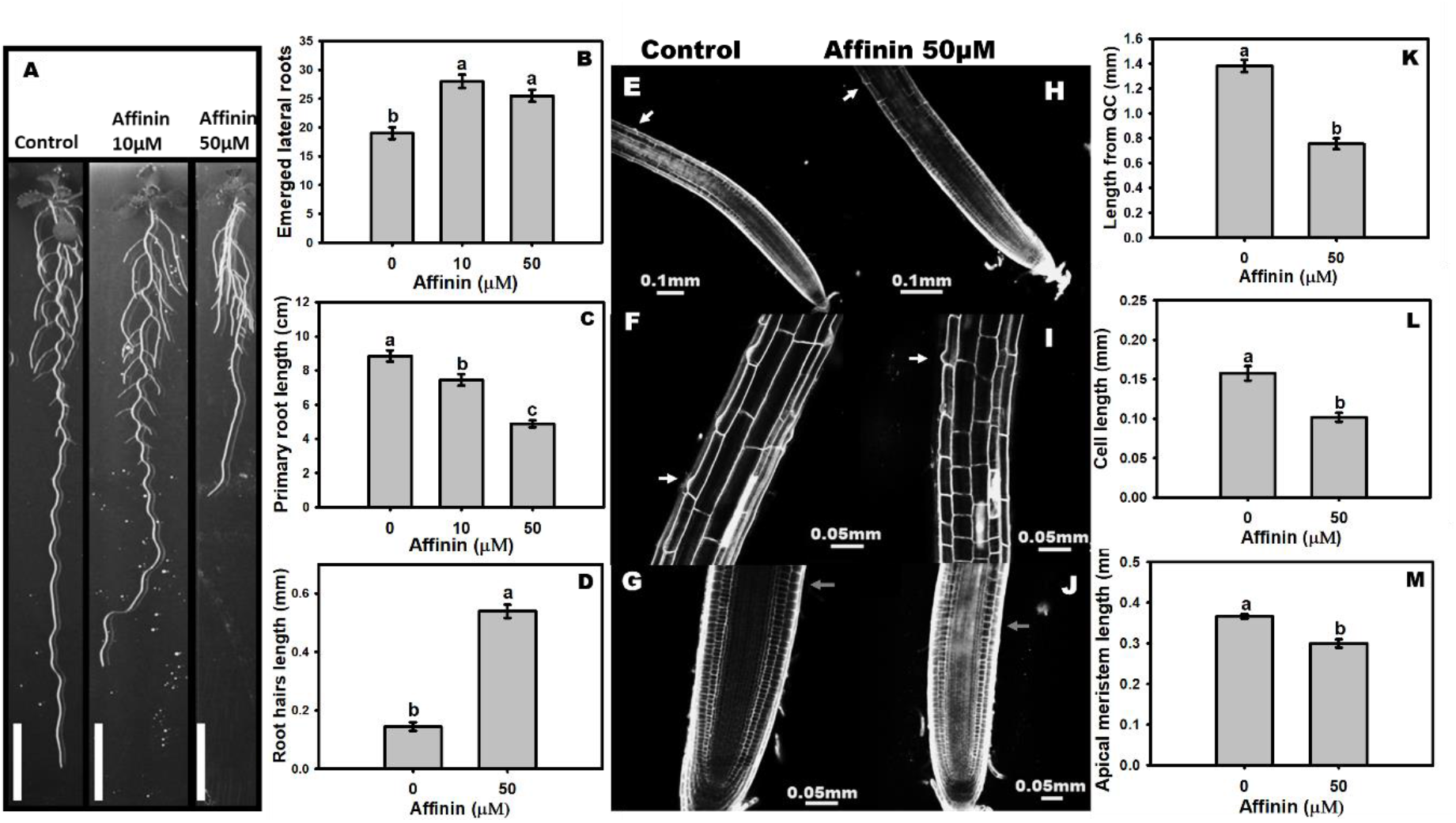
Effect of affinin on root architecture in *Arabidopsis* Col-0 accession. (A) Plants grown on affinin treatments (Bars=1cm); (B) primary root length (*n*=10); (C) number of emerged lateral roots (*n*=10) and (D) root hair length (*n*=10). (E, H) confocal images of the primary root from tip to first root hair bulge (Bars=0.1mm), (F, I) confocal images of cells from the elongation-differentiation zone (Bars=0.05mm); (G, J) confocal images of apical meristem size. White arrows indicate root hair bulge and grey arrows shows the transition zone (Bars=0.05mm). (K) Shows the mean distance from QC to the first root hair bulge (*n*=5), (L) is the mean value of cell length in the elongation/differentiation zone (*n*=3) and (M) shows the mean value of the apical meristem length (*n*=5). Data was analysed with one-way ANOVA followed by a Tukey test using the InfoStat software (www.infostat.com.ar) Different lower-case letters are used to indicate means that differ significantly (*P* ≤ 0.05). Micrographs were adjusted with the same settings; Colour saturation: 0%, Brightness: 40% and Contrast: 40%

### Affinin alters H_2_O_2_ accumulation in root meristems and lateral root emergence sites

*Arabidopsis* transcriptional signatures (Méndez-Bravo *et al*. 2011) suggest that *N-*isobutyl decanamide modulates ROS signalling, which is involved in root development (Passardi *et al*. 2006; Manzano *et al*. 2014), prompting us to assay for affinin-induced ROS responses in *Arabidopsis* roots. DAB staining was utilized to visualize affinin-induced changes in H_2_O_2_ accumulation *in situ*. Staining with DAB forms a brown-reddish precipitate in the presence of H_2_O_2_ and peroxidase activity. In the root meristematic zone, the deposition of DAB stain was reduced by treatment with 50 µM affinin, but no change in DAB staining was seen in the elongation or maturation zones (Fig. 2A). O_2_^•-^ accumulation detected by NBT staining remained unchanged (Fig. 2B). DAB staining was significantly higher in the lateral root primordia emergence sites of roots treated with 50 µM affinin but reduced in the apical meristem (Fig. 2A-F). These results suggest that H_2_O_2_ is involved in affinin-induced changes in lateral root emergence and root apical meristem development leading to an increase in emerged lateral roots and a reduction of primary root length in response to the affinin treatments. These results demonstrate a correlation between sites of H_2_O_2_ accumulation and developmental responses to affinin, prompting further exploration of the role of ROS in these processes.

**Figure. 2.**
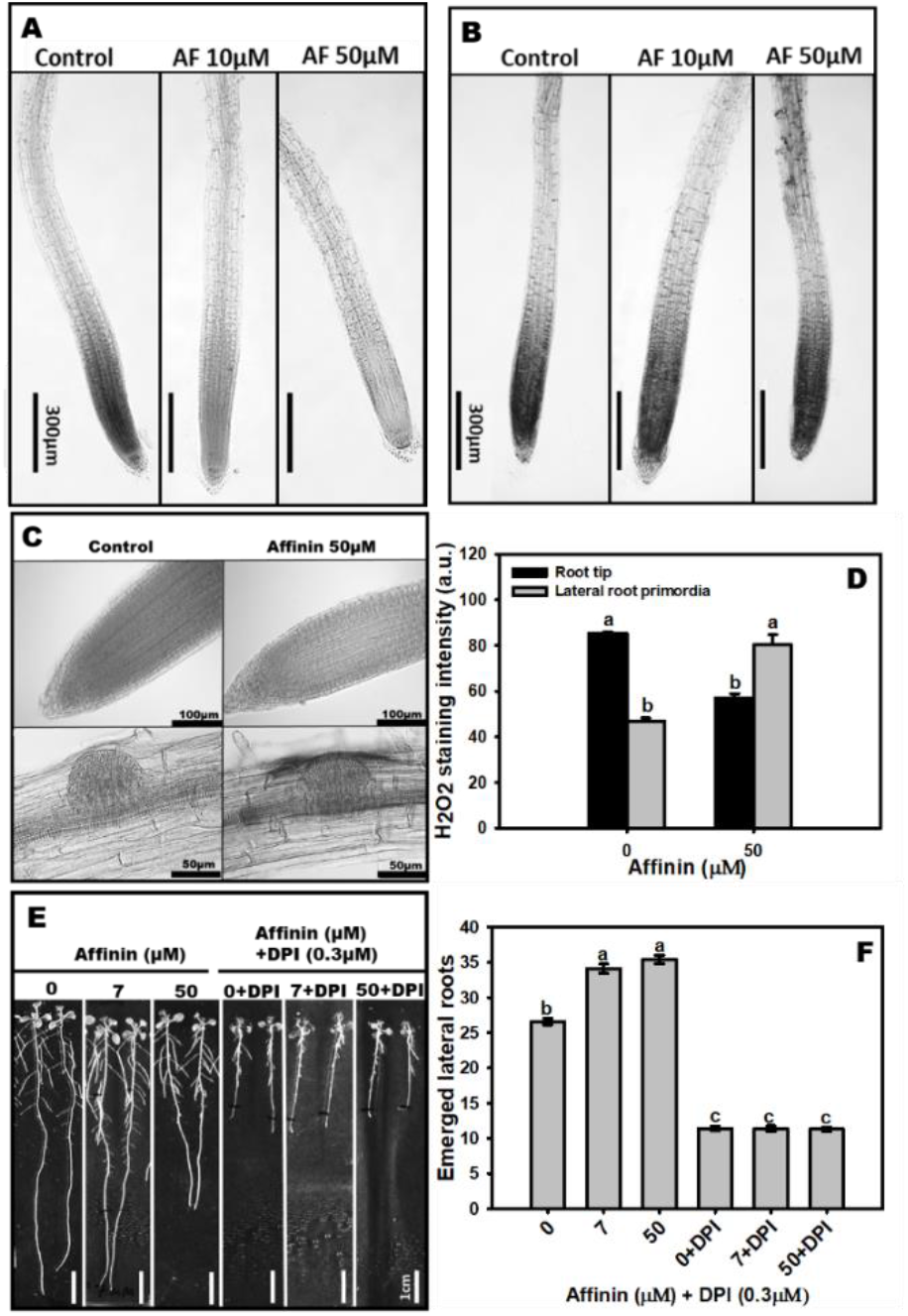
Affinin induced root-developmental changes are mediated by ROS. (A) Detection of endogenous H_2_O_2_ accumulation with 3, 3’-diaminobenzidine (DAB) staining and (B) nitroblue tetrazolium (NBT) staining, visualizing superoxide (O_2_^•-^) accumulation of primary roots treated with affinin (Bars=300µm). (C) Visualization of *in situ* accumulation of endogenous H_2_O_2_ using DAB staining in root apical meristem (Bars=100µm) and lateral root emerging sites (Bars=50µm) of Col-0 wild type plants. Data shown in (D) represent the mean ± SE of H_2_O_2_ staining intensity measured in (C). (E) Affinin-induced *Arabidopsis* (Col-0) lateral rooting is sensitive to synthetic inhibitor of flavoenzymes, diphenyleneiodonium (DPI), used here to target NAD(P)H oxidases (Bars=1cm). Data shown in (F) represent the mean ± SE of emerged lateral root number measured in (E). Data in (D) was analysed with one-way ANOVA (*n*=10) and a Tukey test using the InfoStat software (www.infostat.com.ar). Data in (F) was analysed with two-way ANOVA (*n*=10) and a Tukey test using Minitab software (https://www.minitab.com/). Different lower-case letters are used to indicate means that differ significantly (*P* ≤ 0.05). Micrographs were adjusted with the same settings; Colour saturation: 0%, Brightness: 0% and Contrast: 20%.

### Affinin-induced *Arabidopsis* lateral rooting is dependent of endogenous H_2_O_2_

To test ROS involvement in affinin-induced changes in root architecture, a pharmacological approach was first employed. The effects of affinin on root development were tested in the presence of diphenyleneiododium (DPI), a NADPH oxidase inhibitor. Plants grown in medium containing DPI plus affinin did not show the characteristic affinin induced phenotypes (Fig. 2E). Specifically, seedlings grown on DPI did not increase the number of lateral roots in the presence of affinin; the number of lateral roots in these seedlings remained the same as it was prior to being transferred. Affinin at 7 µM increased the primary root length a response reported by Ramírez-Chávez *et al*. (2004), in contrast primary root growth was totally inhibited by DPI (Fig. 2H-I). These results suggest that endogenous ROS is required for affinin-induced lateral root growth and for primary root development, prompting us to use reverse genetics to define the sources of ROS that are involved in mediating these root architecture changes.

### PRX34-mediated H_2_O_2_ production in affinin-induced responses

The cell wall associated class III peroxidases (PRXs) have been implicated in lateral root formation and regulation of root tip growth (Passardi *et al*. 2006; Manzano *et al*. 2014). The *prx34* mutant (Bindschedler *et al*., 2006) was assayed for affinin-induced root structure alterations and associated DAB stain accumulation. The *prx34* mutant exhibited enhanced inhibition of primary root growth in response to high affinin concentration (Fig. 3A), which was associated with a drop in DAB staining intensity in the meristematic zone (Fig. 3C, D), suggesting that modulation of ROS signalling by PRX34 is implicated in the regulation of meristematic processes by affinin. The number of emerged lateral roots in *prx34* was not increased by affinin treatment (Fig. 3B), while the enhanced DAB staining seen in affinin treated wild type Col-0 plants was not observed in the lateral root emergence sites of *prx34* roots (Fig. 3C, D). These results support that PRX34 was required for alkamide-induced signalling leading to lateral root emergence and suggest that the ROS produced or modulated by PRX34 is involved in this process.

**Figure. 3.**
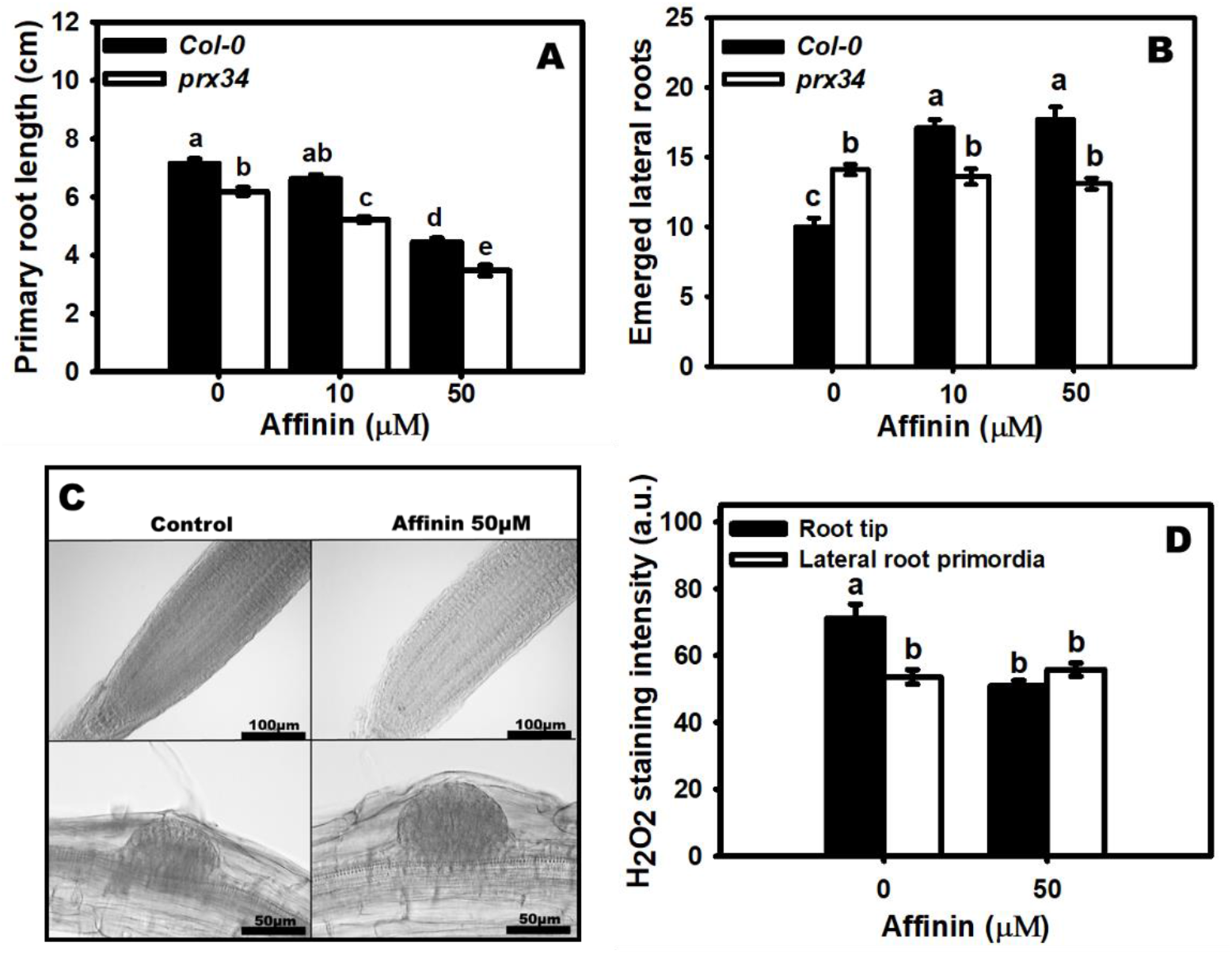
Effect of affinin on root system architecture in the class III peroxidase mutant *Arabidopsis peroxidase34* (*prx34*). (A) Represents the mean ± SE of primary root length (*n*=10) while (B) is the number of emerged lateral roots (*n*=10) of plants grown in different affinin treatments. (C) Visualization of *in situ* accumulation of endogenous H_2_O_2_ using 3, 3’-diaminobenzidine (DAB) staining in root apical meristem (Bars=100µm) and lateral root emerging sites (Bars=50µm) of *prx34* mutant plants. Data shown in (D) represent the mean ± SE of H_2_O_2_ staining intensity measured in (C). Data was analysed with two-way ANOVA (n=10) and a Tukey test using Minitab software (https://www.minitab.com/). Different lower-case letters are used to indicate means that differ significantly (*P* ≤ 0.05). Micrographs were adjusted with the same settings; Colour saturation: 0%, Brightness: 0% and Contrast: 20%.

### NADPH-oxidase mediated ROS production in affinin-induced developmental changes

We next sought to further define the sources of affinin-induced ROS. To explore the contribution of ROS signalling from the RBOH NADPH oxidases in responses to affinin, the root system architecture of wild type plants and the *rbohC, rbohD*, and *rbohF* single mutants were compared following growth in the presence of different affinin concentrations. In medium lacking affinin, all *rboh* mutants exhibited reduced primary root growth as compared to wild type (Fig. 4A) and reduced DAB stain deposition was apparent in the root tips of *rbohC* and *rbohD*, but not *rbohF* under control conditions (Fig. 4C). Primary roots treated with affinin in all mutants showed behaviour similar to WT, however, the *rbohC, rbohD*, and *rbohF* mutants had enhanced growth inhibition at 50µM affinin (Fig. 4A). DAB staining of the apical meristems (Fig. 4C) showed that affinin at 50 µM reduced the H_2_O_2_ accumulation in WT, *rbohC* and *rbohD* but no decrease was observed in *rbohF* (Fig. 4C,E). These results suggest that primary root growth inhibition caused by affinin could be mediated via attenuation of ROS accumulation sourced from RBOHC and RBOHD, but not RBOHF. The *rbohC* and *rbohD* mutants lacked a significant affinin-induced increase in emerged lateral root number (Fig. 4B). DAB staining at lateral roots emergence sites was not enhanced in both mutants (*rbohC* and *rbohD*) treated with 50 µM affinin while in *rbohF* staining was higher in response to 50 µM affinin (Fig. 4D, F) suggesting that RBOHC and RBOHD could be involved in promotion of lateral root emergence via alkamide-induced ROS accumulation.

**Figure. 4.**
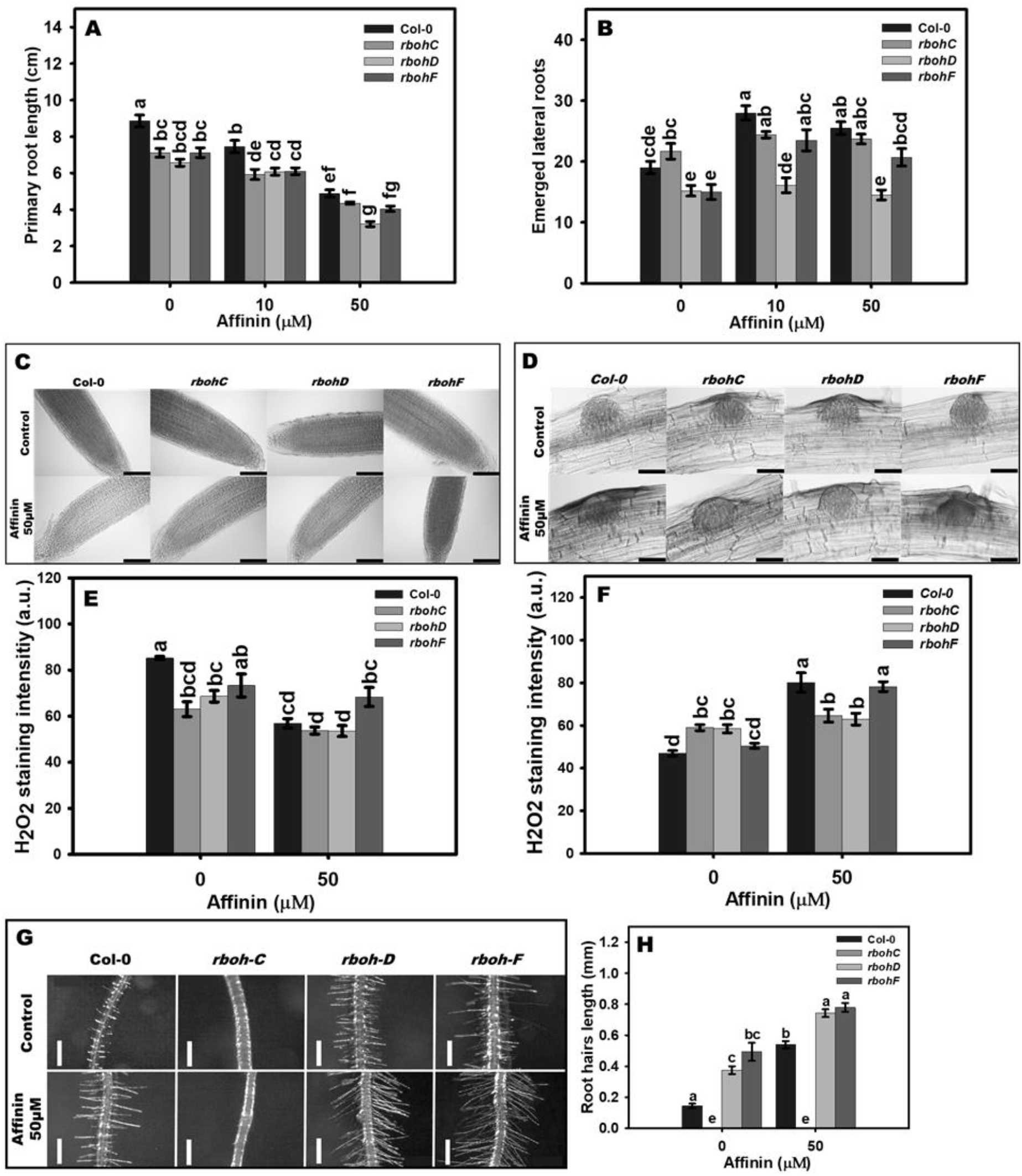
RESPIRATORY BURST OXIDASE HOMOLOG (RBOH) mediated ROS production is involved in root developmental changes induced by affinin. Data shown in (A) represent the mean ± SE of primary root length (*n*=10) while data shown in (B) represent the mean ± SE of emerged lateral roots (*n*=10). (C) DAB staining of root tips and lateral root primordia surrounding area (D). Data in (E) represent the mean ± SE of H_2_O_2_ staining intensity in root tips (*n*=10) and (F) in lateral root primordia surrounding area (*n*=10). Photographs in (G) shown the representative morphology of root hairs, while data shown in (H) represent the mean ± SE of root hair length measured in (G) (*n*=10). Data was analysed with two-way ANOVA (*n*=10) and a Tukey test using Minitab software (https://www.minitab.com/). Different lower-case letters are used to indicate means that differ significantly (*P* ≤ 0.05). Micrographs were adjusted with the same settings; Colour saturation: 0%, Brightness: 0% and Contrast: 20%.

ROS produced by RBOHC is required for cell expansion in root hair growth and characteristically the *rbohC* mutant has greatly reduced root hair length (Foreman *et al*. 2003). Intriguingly, root hair length was unchanged upon affinin-treatment of the *rbohC* mutant indicating that ROS produced by this NADPH-oxidase is necessary for affinin-induced enhanced root hair expansion (Fig. 4G, H). The *rbohD* and *rbohF* mutants had significantly longer root hairs under affinin-treatment (Fig. 4G, H) suggesting that these ROS sources had a minor role as negative regulators of this process. These results highlight the importance of *RBOHC-*mediated ROS production for enhanced root hair expansion.

### G-protein signalling in the *Arabidopsis* affinin response

To test if the heterotrimeric (ht) G-protein subunits, G_α_, G_β_ and G_γ,_ are involved in *Arabidopsis* response to affinin, affinin-induced root phenotypes of *gpa1*-2, *agb1*-2, and *agg2*-1 mutant plants were examined. In all genotypes primary root length was reduced by 50 µM affinin treatments and in *agb1-2* this effect was slightly but significantly enhanced suggesting AGB2 may act as a negative regulator of this process (Fig. 5A). The *agb1-2 and agg2-1* mutants exhibited and increased number of lateral roots under control conditions, indicating that they negatively regulate lateral root formation (Fig. 5B). Interestingly, *agb1-2* exhibited a decrease in the number of lateral roots under affinin treatment compared to control (0µM affinin), while *agg2-2* remained equal in affinin treatment and control, and *gpa1-2* had a slight but not significant increase, suggesting that all these subunits could be required for enhanced lateral root emergence in response to affinin (Fig.5B). In plants G_α_ proteins tend toward their active GTP bound state (Urano and Jones, 2014) and the seven transmembrane domains (7TMD) containing RGS1 protein negatively regulates GPA1-mediated signalling through its GTPase accelerating protein (GAP) activity (Liang *et al*. 2018). The *rgs1-*2 mutant was also tested and had a reduced primary root length under control conditions (Fig. 5C) indicating RGS2 is required for full primary root growth. Affinin treatment reduced primary root length to a similar extent in both Col-0 WT and *rgs1*-2 (Fig. 5C) suggesting RGS1 is not involved in affinin-induced response on primary root growth. Lateral root emergence was the same in WT and *rgs1*-2 under control conditions, but the affinin-induced increase was enhanced in *rgs1*-2, suggesting RGS1 negatively regulates this response (Fig. 5D). Together, these data implicate multiple htG-protein subunits in the regulation of the affinin response.

**Figure 5.**
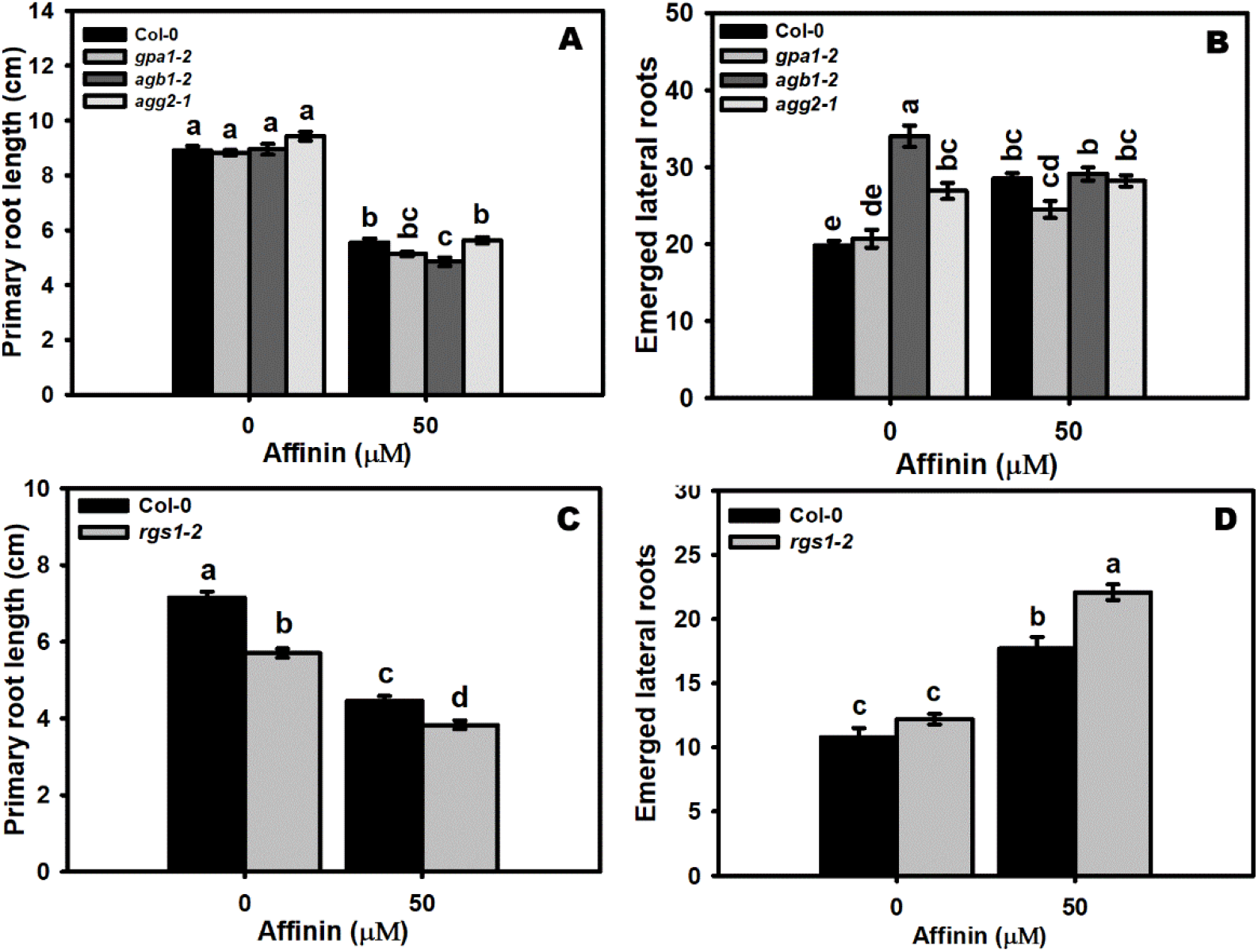
Effect of affinin on root growth of *Arabidopsis* heterotrimeric G-protein subunit mutants. Data shown in (A) represent the mean ± SE of primary root length (*n*=10) and (B) the emerged lateral roots number (*n*=10) from heterotrimeric G-protein mutants while (C) represent the mean ± SE of primary root length (*n*=10) and (D) the emerged lateral roots number (*n*=10) from the regulator of G-protein mutant. Data was analysed with two-way ANOVA (n=10) and a Tukey test using Minitab software (https://www.minitab.com/). Different lower-case letters are used to indicate means that differ significantly (*P* ≤ 0.05).

### Extracellular pH and affinin-induced responses

To test if affinin causes changes in apoplastic pH, an affinin pulse for 24 hours (see Materials and Methods) was applied to seven-day-old *Arabidopsis* roots and changes in the pH in surrounding growth medium was monitored with the indicator bromocresol purple. The results show that affinin treatment induced extracellular acidification in the root elongation zone (Fig. 6A, B). To test the potential functional consequence of these extracellular pH changes, the extracellular pH was stabilized using a medium containing 2-*(N*-morpholino) ethanesulfonic acid (MES) buffer (pH 5.7), which has an active buffering capacity in the relevant range of pH 5.5 - 6.7. In buffered medium, affinin treatment exhibited typical inhibition of primary root growth; however, the number of emerged lateral roots did not increase in response to affinin (Fig. 6C-E). Further, in line with previous studies (Kagenishi *et al*. 2016), root hair growth was inhibited in control plants on buffered medium without affinin; however, root hair length still increased upon affinin treatment on MES buffered medium (Fig. 6F, G). We conclude that the affinin-induced changes in lateral root emergence, but not root hair length, were dependent on a change in extracellular pH.

**Figure 6.**
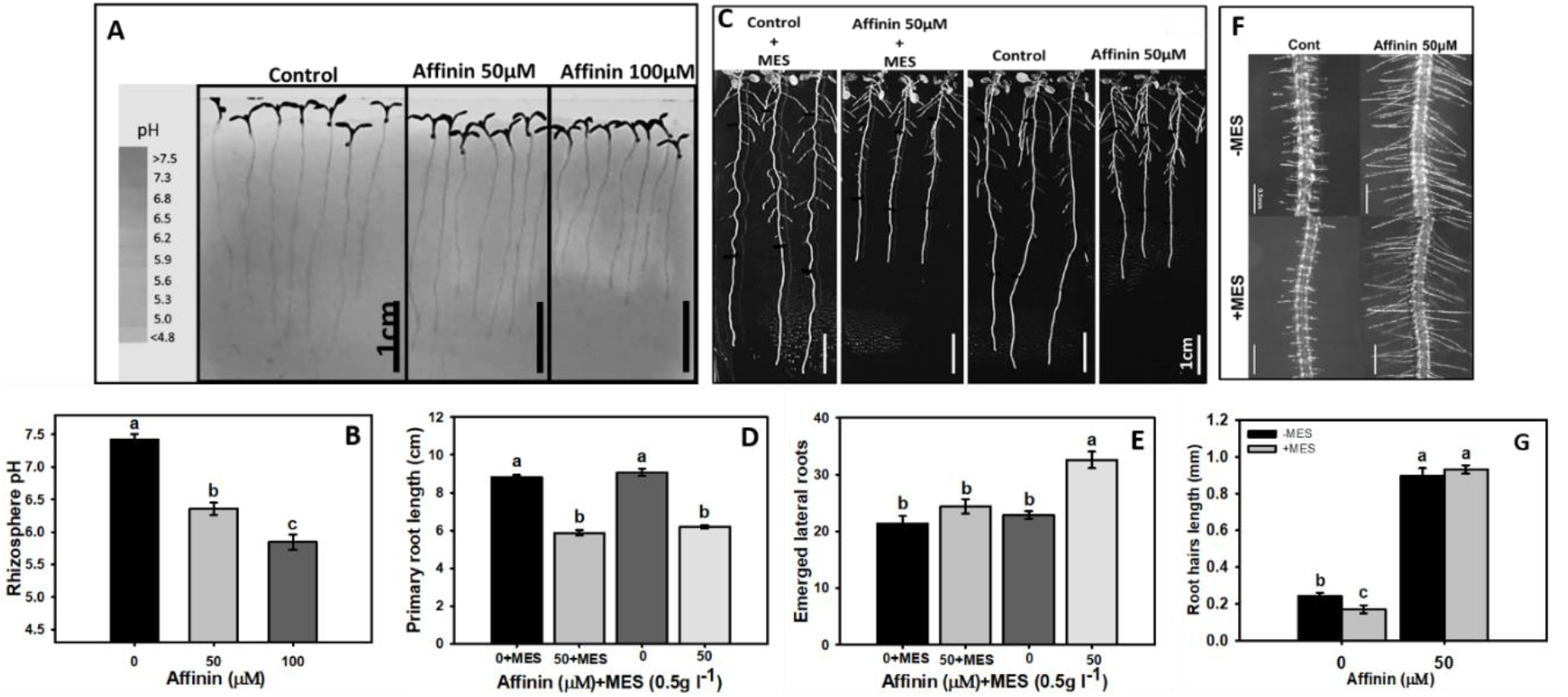
Involvement of extracellular pH change in affinin-induced developmental responses. (A) Rhizosphere acidification in the maturation zone of *Arabidopsis* roots in response to different affinin treatments (Bars=1cm); the scale on the left is a pH reference scale in the agarose without plants. Data shown in (B) represent the mean ± SE of medium pH measured in a region of interest (ROI) near the roots of seedlings (*n*=7). (C) Photographs of full seedlings (Bars=1cm) and (F) root hairs (Bars=0.5mm) of plants treated with or without affinin grown in a medium containing or lacking MES buffer (0.5g L^-1^). Data shown represent the mean ± SE of primary root length (D), emerged lateral roots (E) and root hair length (G) (*n*=10). Data was analysed with two-way ANOVA and a Tukey test using Minitab software (https://www.minitab.com/). Different lower-case letters are used to indicate means that differ significantly (*P* ≤ 0.05). Micrographs were adjusted with the same settings; Colour saturation: 0%, Brightness: 0% and Contrast: 20%.

## Discussion

### The role of ROS in alkamide-induced processes

Decanamide-induced transcriptional reprogramming previously observed in *Arabidopsis* (Mendez-Bravo et al. 2011) suggests ROS signalling may mediate alkamide-induced processes. The current study utilizes a genetic approach with the previously established model system (Ramírez-Chávez *et al*. 2004; Méndez-Bravo *et al*. 2010) of alkamide-induced alterations to *Arabidopsis* root system architecture to demonstrate that these developmental changes are associated with changes in ROS accumulation, as monitored using NBT to visualize O_2_^•-^ production and DAB for H_2_O_2_ (Figs. 2-4). Additionally, this study builds upon the previous observation of affinin-induced H_2_O_2_ accumulation in *Arabidopsis* leaves (Méndez-Bravo *et al*. 2011) by expanding findings to root tissues undergoing developmental changes. We found that 50 µM affinin induced the accumulation of H_2_O_2_ in the peripheral cells surrounding the lateral root primordia (Fig. 2C, D), which coincided with an increased number of emerged lateral roots (Fig.1A, B). These data are consistent with the known signalling role of H_2_O_2_ in lateral root development as previously reported (Passardi *et* al. 2006; Manzano *et al*. 2014; Tsukagoshi *et al*. 2016; Orman-Ligeza *et al*. 2016) and suggest that H_2_O_2_ is involved in root response to alkamides by acting as a signalling intermediate. Further, these results are consistent with the ability of ROS to increase lateral root numbers due to the activation of lateral root pre-branch sites and lateral root primordia. Specifically, ROS are deposited in the apoplast of these cells during lateral root emergence (Orman-Ligeza *et al*. 2016).

Moreover, our results indicate that affinin attenuated growth of the primary root in a dose dependent manner (Fig. 1A, C), which was accompanied by attenuated DAB staining intensity specifically in the root meristematic region (Fig. 2A, C, D). Root growth is controlled by both the rate of cell division in the meristematic zone and the degree of cell expansion in the elongation zone (Beemster and Baskin, 1998; Tsukagoshi *et al*. 2010). Increased root meristem size is correlated with an acceleration of root growth and is a result of increased rates of cell division and the delay of cell expansion (Ubeda-Tomas *et al*. 2009; Tsukagoshi *et al*. 2010). It has been demonstrated that H_2_O_2_ and the O_2_^•^ both have distinct accumulation zones and distinct roles in the growing *Arabidopsis* root (Dunand *et al*. 2007). Our results demonstrate that affinin altered the accumulation of H_2_O_2_, but not O_2_^•^, in the meristem (Fig. 2A, B). Affinin treatment reduced the cell size of the root elongation-differentiation zone (Fig. 1E-M) without altering H_2_O_2_ and O_2_^•-^ accumulation in this root zone (Fig. 2A, B) indicating that affinin can modulate a defined developmental program. This profile of changes is consistent with the action of peroxidases, which are known to modulate primary root growth (Passardi *et al*. 2006). These results agree with the findings that affinin alter meristematic root growth and the expression of *CycB1*gene (Ramírez-Chávez *et al*., 2004).

We also present evidence supporting the involvement of extracellular acidification in response to affinin. Ion fluxes are a common event that often works in concert with ROS signalling in stress and developmental responses. It is known that changes in apoplastic pH are likely to modulate the activity of several regulatory elements such as cell wall proteins, such as expansins (Cosgrove *et al*. 2002) and pectin methylesterases (Sherrier and Vandenbasch, 1994, Micheli, 2001); plasma membrane proteins, such as pH-sensitive potassium channels (Ilan *et al*. 1996, Hartje *et al*. 2001); and is functionally related with the regulation of root hair growth (Monshausen *et al*. 2007).

Beyond correlating ROS accumulation with affinin-induced changes in root system architecture, pharmacological treatments and mutant plants with compromised ROS pathways (Figs. 3-5) were utilized to reveal a requirement for these ROS-signalling pathways for a subset of affinin-induced developmental responses. Based on this pharmacological and genetic evidence, we conclude that ROS-signalling was involved in affinin-induced responses. This work also defined the sources of various affinin-responses, as further discussed below.

The production of ROS in the apoplast depends on several classes of enzymes, most notably, NADPH-oxidases and class III peroxidases. Several NADPH-oxidases are known ROS sources that mediate cell expansion and determines root system architecture (Torres, 2010). These are known in *Arabidopsis* as RESPIRATORY BURST OXIDASE HOMOLOGS (RBOHs) and act specifically in lateral root emergence, and root hair cell expansion (Foreman *et al*. 2003; Orman-Ligeza *et al*. 2016), while class III peroxidases, such as *AtPrx34*, have been associated with an active role in root cell elongation (Sagi and Fluhr, 2006; Passardi *et al*., 2006).

To test if these ROS sources are involved in affinin-response, several available *Arabidopsis* mutants were utilized, including *prx34*, which is defective in a class III peroxidase, and the NADPH-oxidase mutants, *rbohC, rbohD*, and *rbohF*. Testing the effect of affinin in the *prx34* mutant, the ROS staining pattern of lateral root primordia and the number of emerged lateral roots were indistinguishable from wild type Col-0 plants (Fig. 3C, D). It has been demonstrated that *PRX* activities are important for lateral root development, especially during lateral root emergence (Manzano *et al*., 2014). The effects of affinin on root system architecture has been previously shown to be independent or downstream of auxin signalling (Méndez-Bravo *et al*., 2010). Accordingly, lateral root emergence due to PRX activity also occurs independent of auxin signalling (Manzano *et al*. 2014). The accumulation of DAB staining in root tips was reduced by affinin 50 µM in *prx34* as in WT. It has been reported that peroxidases are involved in the regulation of primary root growth (Tsukagoshi *et al*. 2010) and PRX34 was previously shown to be required as a component of the oxidative burst in response to some pathogens (Bindschedler *et al*., 2006). Our results strongly suggest that PRX34 is not required for affinin-induced inhibition of primary root growth, suggesting other PRXs may regulate this process. The reduced accumulation of DAB staining in root meristems upon 50 µM affinin treatment suggests that ROS homeostasis is somehow disrupted due to affinin treatment leading to an alteration in plant cell cycle. This result is supported by the fact that affinin at high concentrations reduced the expression of genes associated with the mitotic cycle, such as *CycB1* (Ramírez-Chávez *et al*., 2004). On this topic, it has been found that cellular ROS signalling oscillations are rapidly transmitted through MAPK pathways inducing MAP activation and affects microtubules dynamics and organization (Livanos et al., 2012).

Using the *respiratory burst oxidase homologues* (*RBOH*) mutants (*rbohC, rbohD* and *rbohF*) these ROS signalling genes were tested for involvement in alkamide induced signalling. This revealed that primary root growth inhibition by affinin was independent of all the loci represented by these mutant lines (Fig.4A), however H_2_O_2_ accumulation as monitored by DAB staining in the apical meristem was reduced in the *rbohC* and *rbohD* mutants while unchanged in *rbohF* (Fig. 4C, E). Interestingly the *rbohD* mutant exhibited a significant inhibition of primary root length in response to affinin compared to the other two mutants and wild type. Taken together, the primary root length and DAB staining results do not support that the ROS produced by *rbohC and rbohF* are involved in affinin-induced response in meristematic activity. However, the impact of affinin on root length in *rbohD* supports the hypothesis that ROS homeostasis could be involved in mediating affinin response in meristematic cells. On the other hand, *rbohC* and *rbohD* failed to increase the number of emerged lateral roots in response to affinin (Fig. 4B) and the DAB staining intensity did not show differences in lateral root primordia between the treatments of these mutants (Fig. 4D, F). These results are consistent with Orman-Ligeza *et al*. (2016), who found that *RBOH*-mediated ROS production facilitates lateral root outgrowth by promoting cell wall remodelling of overlying parental tissues. Indeed, the diverse transcription patterns suggest that *RBOHs* function in broad aspects of growth and physiological response (Sagi and Fluhr, 2006). In contrast, *rbohF* response in terms of emerged lateral roots was similar to the wild type of response induced by affinin and the DAB staining showed similar results (Fig. 4D, F).

The effect of affinin on root hair growth has been demonstrated (Ramírez-Chávez, *et al*. 2004) and corroborated (Fig. 4G). Interestingly, *rbohC* was the only mutant that did not respond to affinin-induced enhanced root hair elongation (Fig. 4G), this result not only demonstrates the importance of ROS in affinin induced signalling but also the specificity of ROS produced by RBOHC on root hair growth. The *rbohD* and *rbohF* mutants exhibited an increase in root hair length higher than the wild type seedlings (Fig.4G), which indicates that these ROS sources could be negative regulators of root hair growth.

In ROS metabolomic studies, the “oxidative stress” signature includes accumulation of several compounds implicated in ascorbate and glutathione synthesis and degradation pathways, and phytohormones such salicylic acid and jasmonic acid (Noctor *et al*., 2016). Additionally, this signature also includes several compounds whose connections to antioxidant metabolism and redox homeostasis are not as obvious. For example, the accumulation of branched chain amino acids induced by ABA treatment, a response that seems to occur reproducibly during redox signalling (Ghassemian *et al*., 2008; Noctor *et al*., 2015). Recently in tomato seedlings it was found that, affinin induced a dose-dependent metabolomic reprogramming that lead to enhanced accumulation of amino acids, organic acids, sugar alcohols, phenolics, and fatty acids (Campos-García and Molina-Torres, 2021), all metabolites that have been associated with marker metabolites in the oxidative stress response.

### The Alkamide–induced developmental program

The details concerning how alkamides affect plant growth remain poorly characterized. Exogenous application of affinin to plants has multiple effects on several plant processes, some of which are similar to responses triggered by some well-known stress related signals. Comparison of affinin-response to these other better characterized signalling pathways may offer insight into affinin-induced signalling.

Biotic and abiotic stresses induce the so-called stress-induced morphogenic response (SIMR), which shares some similarities with the alkamide response. NO is an intermediate in both SIMR (Potters et al., 2009) and alkamide signalling mediating alteration in root system architecture in *Arabidopsis* (Méndez-Bravo *et al*. 2010). Altered root branching, inhibition of cell elongation, as well as changes in apoplastic pH, ROS, and redox signalling, are all common to alkamide-response and SIMR (Potters et al., 2007, Potters et al., 2009; Tongnetti et al., 2012) It is possible that alkamide-induced stress results in SIMR. However, auxin features prominently in the regulation of SIMR and it was demonstrated that alkamides act via an auxin-independent signalling pathway (Ramírez-Chávez *et al*. 2004). Nonetheless, the possibility that some alkamide-induced responses are related to SIMR may be worth further consideration.

Although auxins are considered the major plant growth-regulating hormones underlying root system architecture adjustment, the discovery of novel signal molecules such as *N*-acyl amides, *N-*acyl ethanolamides (NAE) and *N-*acyl homoserine lactones (AHLs) has shed light on the intricate signalling networks that trigger root system architecture modifications and physiological responses (Ramírez-Chávez *et al*., 2004; Méndez-Bravo *et al*., 2010; Coulon *et al*., 2012; Schikora *et al*., 2016). It has been demonstrated that NAEs and AHLs, compounds that are structurally related to alkamides, have a wide range of effects in plant development (Table 1), some of which overlap with affinin-induced responses. Under *in vitro* conditions, affinin shows a trend towards increasing FAAH’s capacity to metabolize NAE 12:0, suggesting that affinin may have some effects on plants acting via this modulation of NAE metabolism. The structural similarity of affinin to NAEs, suggests that affinin might directly influence FAAH activity in plants, but this will also require further investigation (Faure *et al*., 2015). Recently, Aziz and Chapman (2020) proposed the hypothesis that FAAH proteins hydrolyse a broader range of lipophilic substrates than previously recognized, including affinin, and consequently play a pivotal role in *N*-acyl amide-mediated plant– microbe interactions, a function beyond the established role for FAAH in seedling development in *Arabidopsis*.

**Table 1.** *N-*acyl amides in plant development.

| Type | Chain length:<br>insaturations | Activity | Reference |
| --- | --- | --- | --- |
| NAE | 12:00 | Flower senescence: Increase activity of SOD, Cat, APX and GR. | Zhang *et al.*, 2007 |
| NAE | 12:00 | Reduction of: primary root growth, secondary roots, root hair formation, apex swelling, cell invaginations of the plasma membrane, increase levels of vesicles at the cell periphery, improper cell walls near the meristematic region, disorganize endomembrane system and alter vesicular trafficking.<br>Increase the expression of ABRE genes | Blancaflor *et al.*, 2003 |
| NAE | 12:0 to 18:3 | Inhibition of PLDα activity | Austin-Brown and Chapman, 2002 |
| NAE | 12:0, 18:3 | Reduction of root elongation rate. | Wang *et al.*, 2010;<br>Blancaflor *et al.*, 2003 |
| NAE | 12:0, 14:0, 18:0, 18:1, 18:3, 20:4 | Plant defense: Inhibit alkalization of the extracellular medium, modulate the ion flux in the plasma membrane and Induce PAL expression. | Tripathy *et al.*, 1999;<br>Tripathy *et al.*, 2003 |
| NAE | 12:0, 18:2 | Interacts with the ABA signaling pathway to elicit secondary dormancy | Blancaflor *et al.*, 2014; Wang *et al.*, 2006 |
| AHL | C4-HSL | Intracellular Ca<sup>2+</sup> elevation | Song *et al.*, 2011 |
| AHL | C4-HSL, C6-HSL, C8-HSL | Increase primary root growth and plant biomass. Resistance against necrotrophic pathogens.<br>Increase SA-levels and defense-gene regulation. | Liu *et al.*, 2012;<br>Schenk *et al.*, 2012;<br>von Rad *et al.*, 2008;<br>Schuhegger *et al.*, 2006 |
| AHL | C6-HSL | Alteration in auxin/cytokinin level and herbivore susceptibility | von Rad *et al.*, 2008;<br>Schenk *et al.*, 2012; |
| AHL | C10-HSL, C12-HSL | Inhibition of primary root growth. | Zhao *et al.*, 2015 |

In mammals, polyunsaturated NAEs bind to specific 7TMD G-protein coupled receptors (GPCRs) and activate the htG-proteins that regulate multiple downstream signalling pathways (Abadji *et al*., 1999; Bosier *et al*., 2010). In plants, htG-protein signalling has been implicated in many processes such as plant growth and development. Plant htG-protein signalling is fundamentally different from mammalians systems; with no canonical 7TMD GPCRs, the presence of novel EXTRA-LARGE G-PROTEINs (XLG1, XLG2, and XLG3), and a single Gα subunit that has a low intrinsic GTPase activity and is self-activating in the absence of the GEF activity of GPCRs, thus tends toward a constitutively activated state (Urano *et al*., 2016). Plant G-protein signalling has been implicated in AHL-signalling. GPA1 was required for AHL-mediated elongation of *Arabidopsis* roots (Liu *et al*., 2012). However, we found that inhibition of primary root length by affinin was independent of GPA1.

In *Arabidopsis* NAE 18:3 induces stress responses, autophagy, senescence, chlorophyll catabolic genes, represses chlorophyll biosynthesis genes, and these responses require an intact htG protein complex (Yan et al., 2020). Specifically, this study demonstrated the NAE 18:3 response required AGB1, XLGs (using the *xlg1 xlg2 xlg3* triple mutant) and AGGs (using the *agg1 agg2 agg3* triple mutant; Yan et al., 2020). The NAE 18:3 response was enhanced in the *gpa1* and *rgs1* mutants. This work also demonstrates that, due to functional redundancy in these gene families, higher order mutants are required to see clear results. Our results also showed complex profile of phenotypes with htG-protein mutants: GPA1, AGG1, and AGB1 were required for, and *rgs1* conferred hypersensitivity to affinin-induced increased lateral root emergence. For affinin-induced primary root elongation, GPA1, AGG1, and RGS1 were not required, but the *agb1* mutant conferred a slight hypersensitivity. Taken together, these results suggest the similarities between affinin and NAEs warrant further exploration. Especially, further studies with additional higher order G-protein mutants will be required to resolve this question.

Interestingly, the *agb1-2* and *agg2-1* mutants exhibit an increased number of lateral roots under control conditions, and it is known that Gβγ-dimer restrains lateral root formation (Chen *et al*., 2006). Moreover, it is reported that the *agb1* mutant has a more expanded root architecture presumably due to an increased cell proliferation and lateral root formation (Urano *et al*., 2016). Our results show that in the *agb1-2* mutant, primary root length and lateral root number were reduced in response to affinin, which is a contrasting response compared to the mutant phenotype. Taken together these results suggest the htG-protein complex mediating plant response to affinin, modulating cell proliferation activity and lateral root emergence. Further studies will be required to clarify the mechanism involved.

## Conclusions

Our results provide clear evidence that ROS are molecular intermediates involved in lateral roots emergence and root hair growth in response to alkamides. This is supported by the known roles of ROS as mediators and activators of structural changes in roots. In addition, we demonstrate that alkamides induced root extracellular pH changes, which had an effect on lateral roots emergence. These results provide us with evidence that alkamide-induced modification of root architecture depends on modifications in extracellular pH and ROS homeostasis in the lateral root emergence zone. Further, we also provided evidence that heterotrimeric G proteins can be mediating affinin-induced signalling. This is supported by evidence that some G protein subunits may been activating ROS synthesis, inducing softening of cell walls and facilitating lateral roots emergence. While, on the other hand, ROS formed specifically by RBOHC induce Ca ^+ 2^ channels hyperpolarization, allowing Ca entry into the cell and activating root hairs elongation. Nevertheless, further studies with additional higher order G-protein mutants will be required to resolve how htG-proteins are involved in alkamide-signaling.

## Acknowledgements

We thank Tuomas Puukko, Airi Lamminmäki, and Leena Grönholm, for excellent technical support, Eveliina Karjalainen for assistance with mutant seed production, Dr. Huitzimengari Campos-García for help with the statistical analysis, Dr. Vincent-Cervantes Bueno for his help with the confocal microscope, and M.Sc. Enrique Ramírez-Chávez for his support in affinin purification and GC-MS analysis. The members of the Plant Stress Metagroup are acknowledged for the helpful comments and discussions during this project.

## Author Contributions

Project conception and planning, TCG, KO, JM-T; experiment design, TCG, KO; experimental work, TCG; writing the manuscript, TCG, KO; editing and approval of manuscript, TCG, KO, JM-T.

## Data availability

The data that support the findings of this study are openly available at Campos-García *et al*. (2021): Alkamides and ROS signalling. figshare. Dataset. https://doi.org/10.6084/m9.figshare.17303615.v1.

